# Spontaneous Resolution in Racemic Solutions of N-trifluoroacetylated *α*-aminoalcohols

**DOI:** 10.1101/507657

**Authors:** Dmitry V. Zlenko, Anatoly M. Zanin, Aleksey A. Skoblin, Vsevolod A. Tverdislov, Sergey V. Stovbun

## Abstract

The spontaneous resolution was observed in the racemic solution of N-trifluoroacetylated *α*-aminoalcohol (TFAAA-6) in CCl4. In against other cases of the conglomerates formation, the TFAAA-6 forms highly anisometric crystalline structures (strings). Herewith, the spontaneous resolution was not observed in the racemic solution of TFAAA-5 in heptane, where the isometric precipitate was formed. The latter was also observed in the TFAAA-5 solutions in heptane with small enantiomeric excess (EE), down to 2%. With that, the homochiral strings formed in the TFAAA-5 solutions in heptane with larger EEs. In this case, the strings formed from the excess of one of the enantiomers remained in solution after precipitation of the racemic residual. This process leads to the enhancement of chiral polarization in systems close to racemic and can explain the chiral purity of the living cell.

## 1. Introduction

The spontaneous resolution was discovered and described by Louis Pasteur in the solutions of sodium-ammonium tartrate [1]. Two types of isometric crystals (left- and right-handed) formed when crystallization was held at the temperature below 27 °C. The solutions obtained from crystals of only one type were optically active, and the direction of the polarized light rotation depended on the crystal type. At the elevated temperature (above 27°C) only one type of the crystals was found, and the solutions of these crystals were not optically active. The effect of spontaneous enantiomers separation by crystallization was described for many other systems [2, 3], and is tightly related to the solubility of the enantiomers [4] and thermodynamics of crystallization [5, 6]. Nevetheless, the problem of the chiral purity of the living cell, as well as the problem of the initial separation of the individual chemical species are two main questions of the life origin on Earth [7, 8].

The chiral symmetry breaking down was described for several cases. First of all, it is an asymmetric synthesis that became possible if the reaction proceeds under a chiral catalyst action. The best examples of such processes are all biochemical reactions catalyzed by proteins. Besides that, the synthesis of the chiral compounds in the presence of chiral catalyst was described for non-living systems [9, 10, 11, 12]. The spontaneous symmetry breaking can occur in complex chemical reactions described by Frank [13] and then observed experimentally [14, 15]. The chirality can amplify in the system of self-replicating units due to the difference in the rates of amplification of the chiral and racemic molecules [16, 7]. The symmetry could break spontaneously during the crystallization process, as it was demonstrated for the NaClO_3_ solutions if crystallization proceeded under stirring [17] or in the presence of some additional components, like glass beads [18, 19, 20]. The crystallization combined with the milling and a racemization processes could yield products of very high chiral asymmetry, up to 100% in ideal conditions [21].

As we have reported earlier, the chiral solutions of N-trifluoroacetylated *α-*aminoalcohols (Fig. 1) gelate due to the formation of the supramolecular strings having a length up to several millimeters and diameter of several microns [22]. Strings are highly-elongated supramolecular fibers that could be considered as a quasi-one-dimensional crystals due to ordered packing of the TFAAA molecules in them. However, the achiral TFAAA or a racemic mixture of chiral TFAAAs do not gelate and form strings. So, the gelation is strongly related to the chirality in this case and it reasonable to propose that the spontaneous resolution effects could also be possible here. According to the X-ray, the strings have a crystalline structure, but their shape and dimensions do not allow to use classical approaches and verify if the strings mixture is a conglomerate or a racemate.

**Figure 1:**
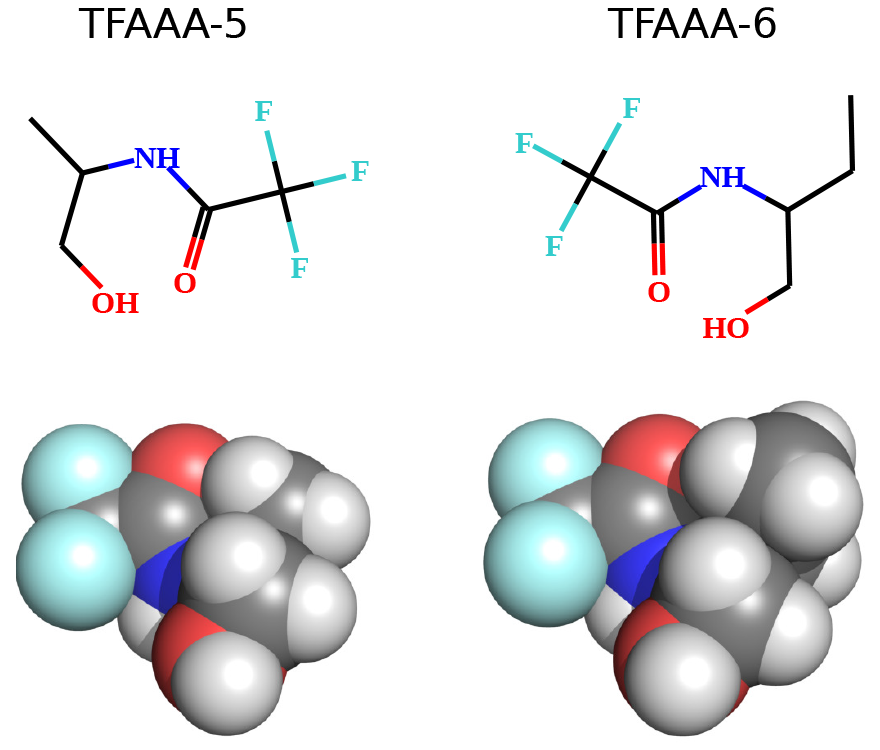
The structural formulas and Van der Waals models of the TFAAA-5 and TFAAA-6 molecules.

In this work, we present the results on the spontaneous resolution of enantiomers in the TFAAA gels containing conglomerates of extremely anisometric crystalline strings. For our knowledge, the conglomerates are usually composed of more or less isometric crystals [1, 23, 5, 4, 19]. Thus, our case of spontaneous resolution is rather unusual and due to anisometry of the strings could be related to the early stages of biological evolution on our planet. Indeed, the elementary strings having the thickness of 1–2 nm and length of at least several microns resembles some of the biological macromolecules, first of all, the DNA double helix. The observed example of the supramolecular self-ordering, when the strings grow in homogeneous solution of small (Mr 200 Da) chiral compounds allows proposing an alternative hypothesis for early stages of chemical evolution considering the self-ordering as a first step and a polymerization as the next one. We also found the self-ordering and chiral strings growth in solutions with rather small (~ 2 %) enantiomeric excess, that could be one of the reasons of why all the chiral biomolecules always present by only one enantiomer.

## 2. Results and discussion

### 2.1. TFAAA-6 gels

The strings were observed microscopically in the dried gels (xerogels) obtained from homochiral as well as from racemic solutions of TFAAA-6 in CCl_4_. The strings were also observed in xerogels of the homochiral TFAAA-6 obtained after heptane evaporation, but was not observed for racemic TFAAA-6 solutions. The morphology of the homochiral and racemic CCl_4_-xerogels was identical (Fig. 2). The length of the strings was also the same in homochiral and racemic xerogels (up to several millimeters). The X-ray diffractograms of the homochiral and racemic gels demonstrated only minor differences (Fig. 3A). The observed differences could be explained by the differencies in the mechanical stress in different specimens. Indeed, the Bragg’s law for the ideal infinite lattice is:

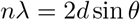

where *n* – is a positive integer, λ – the wavelength, *d* – is the interplanar distance, and *θ* – ia a scattering angle. Assuming *n* = 1, for two lattices having interplanar distances of *d* and *d+δd* (*δd* ≪ 1), for the deviation of the scattering angle *δθ* the Bragg’s law gives:

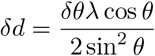

**Figure 2:**
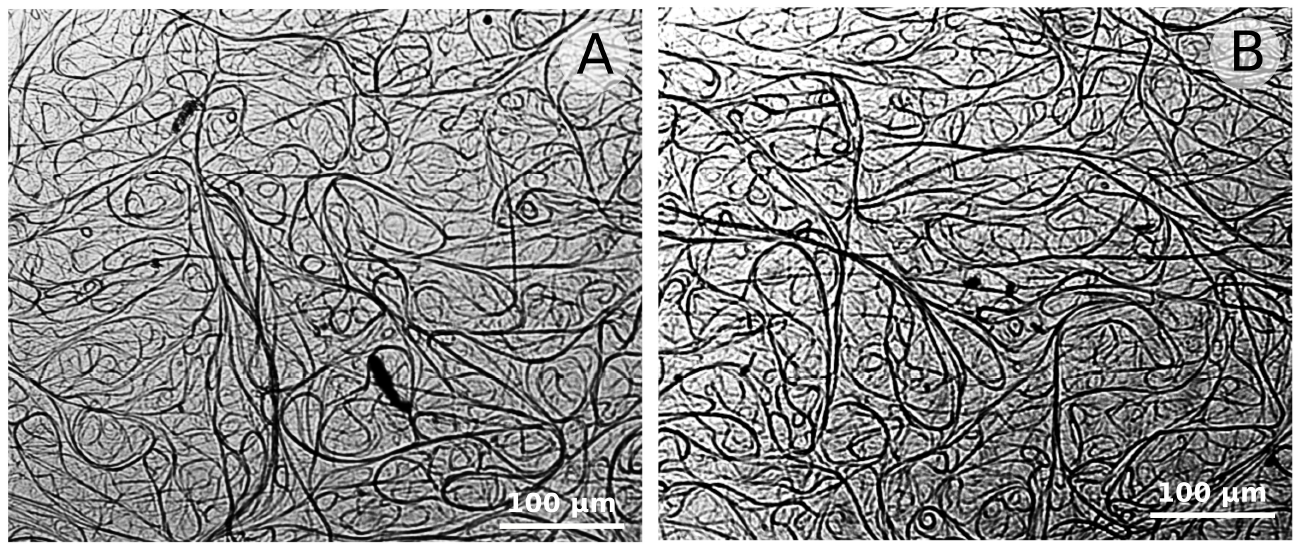
Morphology of homochiral (TFAAA-6D, A) and racemic (TFAAA-6L/6D, B) xerogels obtained after evaporation of the CCl4. Total TFAAA-6 concentration 10 mg/ml.

**Figure 3:**
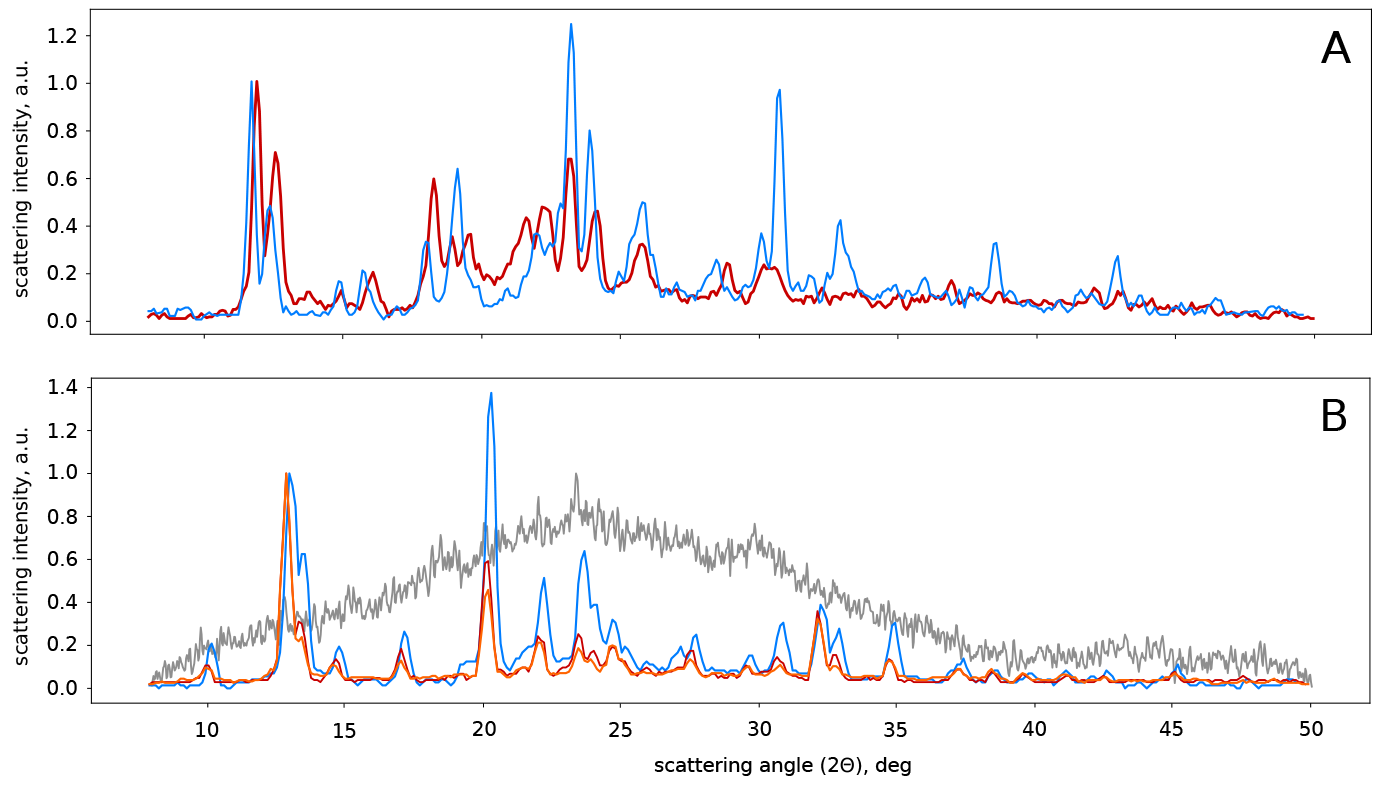
Normalized at 2Θ = 12° X-ray diffractograms of the TFAAA xerogels. A. Homochiral (TFAA-6D, red) and racemic (TFAAA-6L/6D, blue) xerogels after CCl_4_ evaporation. B. Homochiral (TFAAA-5D – red and TFAAA-5L – orange) and heterochiral (TFAAA-5L/5D = 7/3), blue) xerogels after heptane evaporation. Gray curve corresponds to the diffractogramm of the amorphous racemic residual of TFAAA-5. Total TFAAAs concentration 10mg/ml.

In our case (Fig. 3B), λ = 1.5 Å, the *δθ* is about 0.2deg (~ 0.003rad), and *θ* is about 20–30 deg. This leads to the *δd* of about 0.01−0.1 Å. The obtained value lay in the range of the thermal fluctuations and could be achieved due to the stress caused by the strings supercoiling. The latter is always takes place in case of the thick strings [22]. The CD spectra of the TFAAA-6*L* and TFAAA-6*D* gels were very close to each other in shape but had different signs (Fig. 4), while the racemic gel did not demonstrate any CD signal.

**Figure 4:**
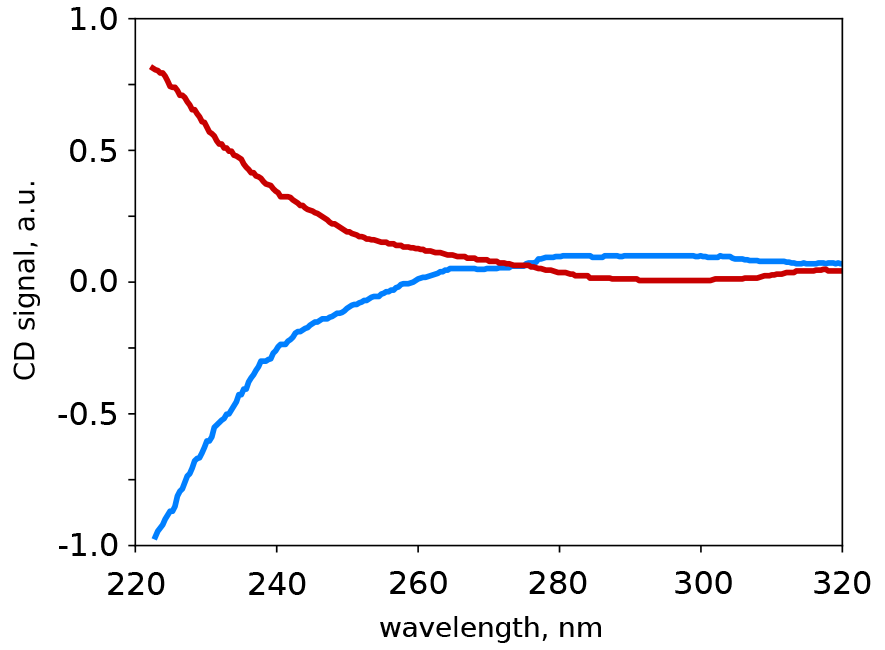
UV CD spectra of TFAAA-6L (red) and TFAAA-6D (blue) gels (10 mg/ml in CCl_4_). Total TFAAA-6 concentration 10 mg/ml.

The identity of the gel scaffolds’ morphology and similarity of their X-ray diffractograms indicates that in the racemic and homochiral gels the strings have the same or very similar molecular structures. However, the CD spectra obtained in the course of the TFAAA solutions cooling showed that the CD signal in this system is defined first of all by supramolecular spiralization rather than by an optical activity of the individual molecules [22]. Since the CD spectra of *L*- and *D*-gels had the same shape and different signs, the crystal lattices of *L* and *D* strings could be transformed into each other by mirroring. In the homochiral gels, the strings’ swirl direction depended on the chirality sign of the TFAAA which was observed microscopically for “vortex-like” structures formed by strings [22]. At the same time, according to the X-ray diffractograms (Fig. 3), the strings in the racemic gels had the same or very similar helical structure as in the chiral gels [22], although both types of the enantiomers were presented in the mixture. Thus, the racemic gels of TFAAA-6 in CCl_4_ contained an equal amount of “left” and “right” elementary strings composed of pure enantiomers. Thus, gelation of the racemic solution of TFAAA-6 in CCl_4_ was accompanied by the spontaneous resolution, and the gel appeared to be a conglomerate. Moreover, instead of the classical more or less isodiametric crystals [1]. TFAAA-6 formed highly anisometric strings that could be considered as a quasi-one-dimensional crystals.

### 2.2. TFAAA-5 gels

The homochiral solutions of TFAAA-5 (both *L* and *D* isomers) in heptane gelated under cooling [22]. In against to the TFAAA-6/CCl_4_ solution, the gel scaffold, in this case, was composed of longer and more straight strings (Fig. 5). The observed difference in strings morphology likely corresponds to the solvent influence, rather than to the TFAAA molecules peculiarities [22]. Indeed, the chiral TFAAA-5 solutions in CCl_4_ also gelated under cooling, and the strings morphology was similar to the observed for the TFAAA-6 in CCl_4_ (Fig. 2). The racemic solution of TFAAA-5 in heptane did not gelate but precipitated under cooling. The continuous drying of the precipitate leaded to the formation of the dense residual. The X-ray diffractogram of this residual indicated its amorphous structure (Fig. 3B, gray curve). The optical microscopy investigations of the residual as well as of the dried supernatant did not reveal any strings or other supramolecular structures. The latter indicated that most of the dissolved TFAAA-5 has settled out. Thus, the spontaneous resolution and enantiomers separation did not occur in the racemic TFAAA-5 solution in heptane.

**Figure 5:**
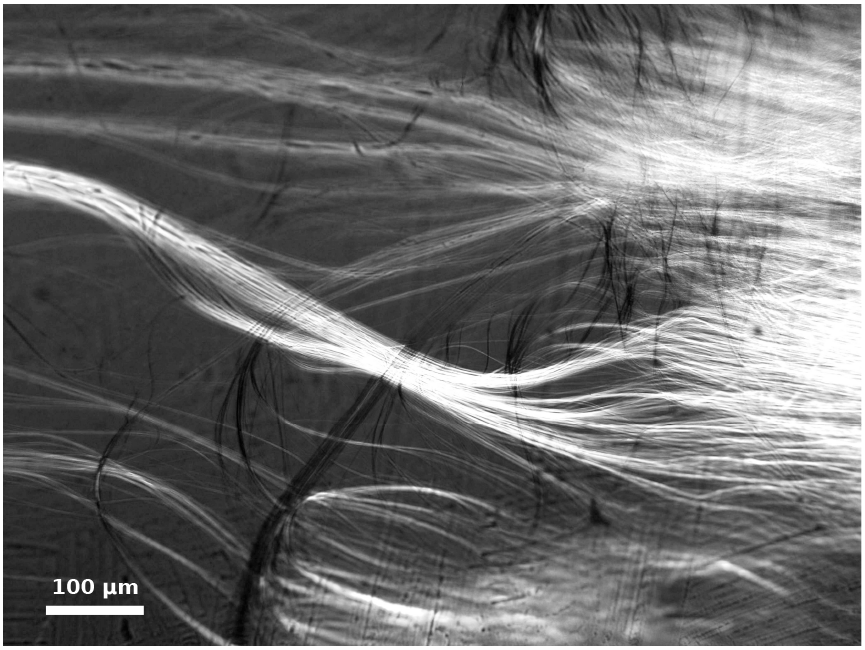
A bundle of strings in xerogel of TFAAA-5L/5D mixture (7:3) after heptane evaporation. Total TFAAA-5 concentration 10 mg/ml.

However, if the TFAAA-5 solution in heptane was not exactly racemic (“heterochiral”) but had some small excess of one of the enantiomers, the solution became capable of gelation under cooling. The strings observed in heterochiral TFAAA-5/heptane xerogels were thin and had a length up to several millimeters, as it was observed in TFAAA-6/CCl_4_. The morphology of the heterochiral TFAAA-5/heptane strings differed from that of TFAAA-6/CCl_4_, as TFAAA-5 formed more straight and branched strings (Fig. 5). Together with the strings, the amorphous precipitate was always observed after cooling of the heterochiral TFAAA-5 solution in heptane, and the amount of this precipitate was always significant.

The X-ray diffractograms of the TFAAA-5*L* and TFAAA-5*D* xerogels were more or less identical (Fig. 3B). The diffractogram of the heterochiral TFAAA-5 xerogel demonstrated some minor but noticeable differences compared to the homochiral one. The observed differences were the same as in case of TFAAA-6 gels by the order of magnitude. Thus, we propose that the reason for this was the same as for the TFAAA-6 gels, i.e. the mechanical stress and deformation of the strings due to their supercoiling.

The gelation ability of the heterochiral TFAAA-5 solutions was determined by the difference between the enantiomer concentrations (*EE*). Moreover, the strings grew and the gel formed if only the *EE* exceeded the threshold concentration of the strings formation (*C*^*^) determined earlier for the homochiral TFAAA-5 solutions in heptane [22]. This could be explained by the isometric precipitation of the most part of the TFAAA-5 molecules, while the dissolved remnant became homochiral (*L* or *D*). This mechanism allows obtaining the solvent of the pure enantiomer from the solution with small initial *EE* (down to 2%). The concentration of the pure enantiomer, in this case, should be approximately equal to *EE*. When it exceeds *C*^*^, the solution gelates, as it was described for homochiral solutions [22]. This assumption defines dependency of the effective string’s formation threshold 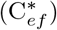 on the “chiral polarization” *δ* = 100% · (*EE/C_tot_*). Indeed, if the racemic fraction precipitates, the real critical concentration of the chiral TFAAA (*C*^*^) in heterochiral solution would be equal to *EE*

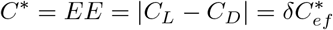

so, the effective critical concentration would have a hyperbolic dependency on the chiral polarization:

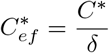

The observed dependency of the effective critical concentration 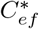 on the chiral polarisation *δ* has a pronounced hyperbolic shape (Fig. 6). The approximation of the obtained dependency by a single-parameter hyperbolic function lead us to the value of the real critical concentration of *C*^*^ ~ 0.04 mg/ml that is very close to the value obtained for the pure solutions of enantiomers [22]. This observation is a very strong argument in favor of the assumption made.

**Figure 6:**
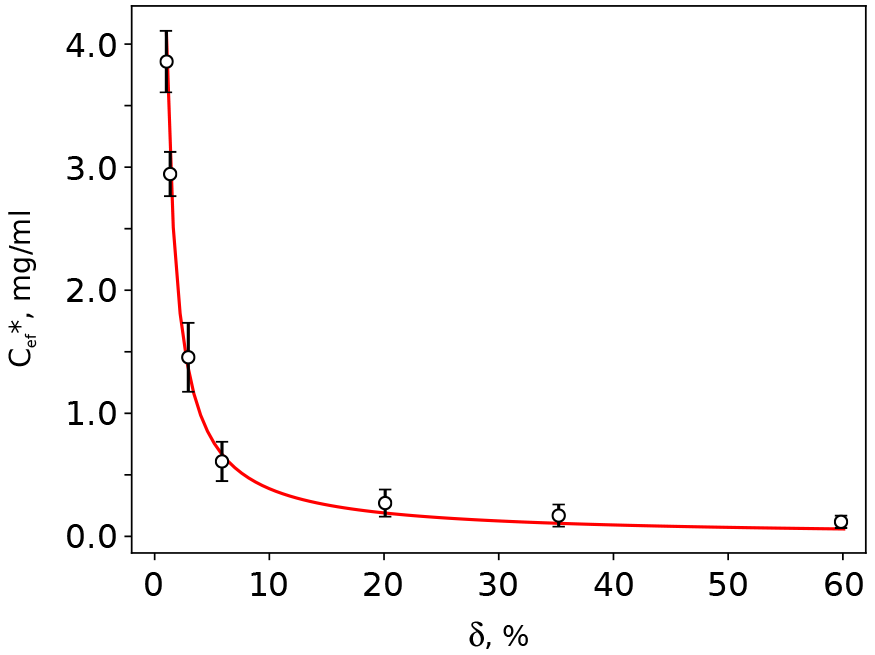
The effective string formation threshold 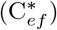 in the heterochiral TFAAA-5 solutions in heptane as a function of the chiral polarisation (*δ* = 100% *EE/C_tot_*). Circles represent the experimental data (standard deviation as an error), while the red line represents a hyperbolic approximation curve: 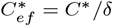, where *C*^*^ ~ 0.04 mg/ml.

## 3. Conclusions

Despite the similarity of the chemical structure and shape of the molecules, racemic solutions of TFAAA-5 and 6 demonstrated the qualitatively different behavior. When cooled, the TFAAA-6 solution in CCl_4_ gelated and undergone the spontaneous resolution. So, the resulting gel appeared to be a conglomerate. On the contrary, the racemic solution of TFAAA-5 in heptane did not gelate, and TFAAA-5 precipitated under cooling. The “heterochiral” TFAAA-5 solutions with some excess of one of the enantiomers gelated in case if *EE* exceeded the critical threshold of gelation (*C*^*^) [22]. The TFAAA-5 and 6 molecules have close structure and shape, as the TFAAA-6 molecule is only one methyl group greater. The observed qualitative difference in their behavior could not be explained by the free energy of the methyl group interaction with the solvent or other TFAAAs. Indeed, the entropy gain caused by combining of two TFAAA molecules is ~ kTln2. The van der Waals energy of the methyl group interaction with the surrounding media lay below 1 kJ/mole [24] that is not enough for the entropy barrier overcome. On the other hand, even a small change of the shape of the molecules can significantly affect the steric interactions, as it is observed, for example, in the case of the substrate-enzyme interactions. As far as the strings are highly ordered supramolecular constructions [22], the stereospecific interaction between the TFAAA molecules seems to be a key point providing for the strings growth. The fact, that we did not observe the resolution of TFAAA-6 in heptane, as well as of the TFAAA-5 in CCl_4_ shows, that the steric interactions between the solvent and solute are also of crucial importance for the supramolecular self-organizing.

There is a clear analogy between the TFAAA strings and molecular helices observed in the biological macromolecules like α-helices in proteins or DNA double helix. In both cases, there are more or less isodiametric molecules that form long and thin helical fibers. The latter have a thickness of about several nanometers that is very close to the reported diameter of the TFAAA elementary strings [22]. The difference is that the biological macromolecules are polymers, while TFAAAs are not, although some biological molecules are known to be capable of the self-ordering into the fiber-like structures [25].

The effect of the heterochiral TFAAA-5 solution gelation provides for the mechanism of enantiomers separation. Indeed, if the racemate of some kind of the molecules easily settles out, while the pure enantiomers are capable of self-assembly, then even a small *EE* can give rise to the formation of chiral supramolecular structures. The difference in the shape of the chiral strings and isometric racemic precipitate lead to their different behavior under any kind of external mechanical influence, such as solvent flow, waves or whatever else. So, the long and chirally pure strings could be separated from more or less isometric racemic precipitate. Moreover, the crystallization itself can provide for separation of individual chemical species [5, 18] from the initial complex mixture [7, 26, 8]. Thus, only capable of self-ordering and stereochemically pure part of the initial population of organic molecules could be separated from all other components of the mixture. In this case, the capability of self-ordering become a key feature providing for the purification, so, it is reasonable to propose that the formation of the helical supramolecular structures was the initial step in early chemical evolution, while the polymerization of the monomers fastened and stabilized them later. This assumption is strongly confirmed by the effect of “enantiomeric cross-inhibition” [16], that implies that the replication process and the related natural selection process could begin only in the stereochemically pure environment.

## 4. Methods

The pure enantiomers (both *L* and *D*) of TFAAA-5 and TFAAA-6 were synthesized as was described earlier [27]. The racemic and chiral solutions of TFAAA-5 in heptane and TFAAA-6 in CCl_4_ were prepared using solvents (ChimMed, Russia) and pure *L* and *D* isomers (Fig. 1). DRON-3 X-ray diffractometer (Russia) with a copper anticathode and a nickel filter (30 kV and 20 mA) was used for the dried TFAAA gels analysis. SKD-2 circular dichroism (CD) spectrometer (Russia) was used for CD spectral analysis of the liquid TFAAAs’ solutions and gels. The gel scaffold morphology in the samples of dried gels was analyzed with MIKMED-6 (LOMO, Russia) optical microscope.

TFAAA gels were obtained as follows: i) sample weight of the TFAAA was dissolved in heated up (60 °C for CCl_4_ and 80 °C for heptane) solvent; ii) after complete dissolution the obtained solution was cooled down that leads to gelation; iii) to obtain the xerogels the sample of the gel or cooled to the room temperature TFAAA solution were incubated for several hours in the air until the complete solvent evaporation. The residual was examined by X-ray diffractometer or optical microscope. The racemic precipitate of TFAAA-5 was dried for a week at room temperature in the air. The resulted dens substance was examined by X-ray diffractometer.

## 5. Acknowledgements

The work was supported by FASO Russia, theme number 45.9, 0082-2014-0011, AAAA-17-117111600093-8.

